# Complementary learning systems within the hippocampus: A neural network modeling approach to reconciling episodic memory with statistical learning

**DOI:** 10.1101/051870

**Authors:** Anna C. Schapiro, Nicholas B. Turk-Browne, Matthew M. Botvinick, Kenneth A. Norman

## Abstract

A growing literature suggests that the hippocampus is critical for the rapid extraction of regularities from the environment. Although this fits with the known role of the hippocampus in rapid learning, it seems at odds with the idea that the hippocampus specializes in memorizing individual episodes. In particular, the Complementary Learning Systems theory argues that there is a computational trade-off between learning the specifics of individual experiences and regularities that hold across those experiences. We asked whether it is possible for the hippocampus to handle both statistical learning and memorization of individual episodes. We exposed a neural network model that instantiates known properties of hippocampal projections and subfields to sequences of items with temporal regularities. We found that the monosynaptic pathway — the pathway connecting entorhinal cortex directly to region CA1 — was able to support statistical learning, while the trisynaptic pathway — connecting entorhinal cortex to CA1 through dentate gyrus and CA3 — learned only individual episodes, with apparent representations of regularities resulting from associative reactivation through recurrence. Thus, in paradigms involving rapid learning, the computational trade-off between learning episodes and regularities may be handled by separate anatomical pathways within the hippocampus itself.

## Introduction

The complementary learning systems (CLS) theory (1) provides a powerful computational framework for understanding the distinct roles that the hippocampus and cortex play in representing memories. It demonstrates the fundamental trade-off between memorizing the specifics of individual experiences (e.g., where I parked my car today), which benefits from separate representations for each experience, and extracting regularities across those experiences (e.g., where parking spaces tend to be open), which benefits from overlapping representations.

To solve this trade-off, CLS posits that the brain recruits different systems: The hippocampus uses a high learning rate and sparse, relatively non-overlapping (*pattern separated*) representations to quickly store memory traces for each recent experience without interference from other similar recent experiences. The hippocampus then slowly teaches these experiences to cortical areas during offline periods, such as sleep. The cortex has a slow learning rate and overlapping representations, which allow it to extract and represent regularities across experiences — over days, months, and years.

Although CLS has been successful in accounting for many empirical findings (2), it has not been used to address an important type of learning: Apart from memorizing individual experiences rapidly and learning regularities across those experiences over long periods of time, we can also learn regularities rapidly — over minutes or hours (3). Critically, there have been several empirical demonstrations that the hippocampus is involved in, and even necessary for, such rapid statistical learning (4–11).

These findings pose a challenge for CLS. In neural network models, regularities are most effectively extracted via overlapping representations, which facilitate generalization between stimuli based on their shared features (1). However, the hippocampus in CLS does not employ overlapping representations — to the contrary, it specializes in minimizing overlap to prevent interference (1, 12). Thus, although the learning rate of the hippocampus is well suited to the timescale of rapid statistical learning, the mechanism by which it can represent regularities is unclear.

To address this puzzle, we ran simulations of statistical learning using a recent version of a hippocampal neural network model that instantiates the episodic learning component of CLS (13). The model explains how known anatomical pathways in the hippocampus and subfield properties might together support episodic memory. It contains hippocampal subfields dentate gyrus (DG), Cornu Ammonis 3 (CA3), and CA1, as hidden layers that learn to map input provided by superficial layers of entorhinal cortex (EC_in_) to output through deep layers of EC (EC_out_). There are two main pathways: the trisynaptic pathway (TSP), EC_in_ → DG → CA3 → CA1, and the monosynaptic pathway (MSP), EC ↔ CA1. The connections between layers within the TSP are sparse, and CA3 and especially DG have high levels of inhibition, making the TSP the engine for pattern separation of episodic memories. This pathway, with the help of strong recurrent connections within CA3, also retrieves previously memorized patterns from partial cues (*pattern completion*). CA1 employs more overlapping representations and a relatively slower learning rate — it is more cortex-like — and has acted as a translator between sparse representations in the TSP and overlapping representations in EC. The role of the MSP has simply been to help the hippocampus communicate with cortex.

We simulated three learning paradigms — pair structure, community structure, and associative inference — that require extracting regularities on the timescale of minutes to hours. All have been linked to the hippocampus, with representations of associated stimuli coming to be represented more similarly (5, 6, 14). We found that the MSP took on a new role when confronted with these kinds of structured input: It learned and represented the regularities. Both representational similarity and recurrent activity dynamics contributed, making contact with a recent computational model of how recurrent activity in the hippocampus can support generalization (15). Combined with the role of the TSP in rapid learning of the specifics of individual experiences, these findings reconcile the trade-off between episodic memory and statistical learning by suggesting that the hippocampus itself contains complementary learning systems.

## Methods

### Model Architecture

We built on a neural network model of the hippocampus that has been successful in accounting for episodic memory phenomena, and incorporates known projections and properties of hippocampal subfields (12, 13, 16). We used a recent implementation (13) in the Emergent simulation environment (17), version 7.0.1. Our project was modified from hip.proj (18), as it was implemented for version 6 of Emergent.

#### Activity dynamics

The model uses “units”, which represent neurons or small populations of neurons, with a rate code (ranging from 0 to 1) corresponding to how fast the neurons are spiking. Units communicate with each other through weights, whose values reflect the efficacy of the synapse(s) between the neurons. A unit’s activity is proportional to the activity of all units connected to it, weighted by the value of each of the connection weights between them. A unit’s activity is also modulated by local inhibition between units within a layer. The inhibition corresponds to the activity of inhibitory interneurons, implemented in these simulations using a set-point inhibitory current with a k-winners-take-all (kWTA) dynamic. See (18, 19) for details and equations.

#### Input/output

Superficial layers of EC (EC_in_) provide input to the hippocampus and deep layers of EC (EC_out_) provide output (Fig. 1). EC_in_ and EC_out_ use orthonormal representations in these simulations — each item in the paradigm was represented by activation of one unit (with the number of units in EC_in_ and EC_out_ varying across paradigms). There is also a separate Input layer, not shown, with the same number of units as EC_in_ and one-to-one connections to EC_in_. Input was clamped in this layer so as to allow EC_in_ to also receive input from EC_out_, completing the “big loop” of the model. To clamp input, the units in the Input layer corresponding to the current and previous stimulus were forced to maintain high activity over the course of the entire trial, while all other units were forced to have no activity. Activity traveled from the Input layer to the stimulus representations in EC_in_, then to the rest of the network. EC_in_ and EC_out_ each had inhibition set so that two units could be active at a time, unless otherwise noted.

**Figure 1.**
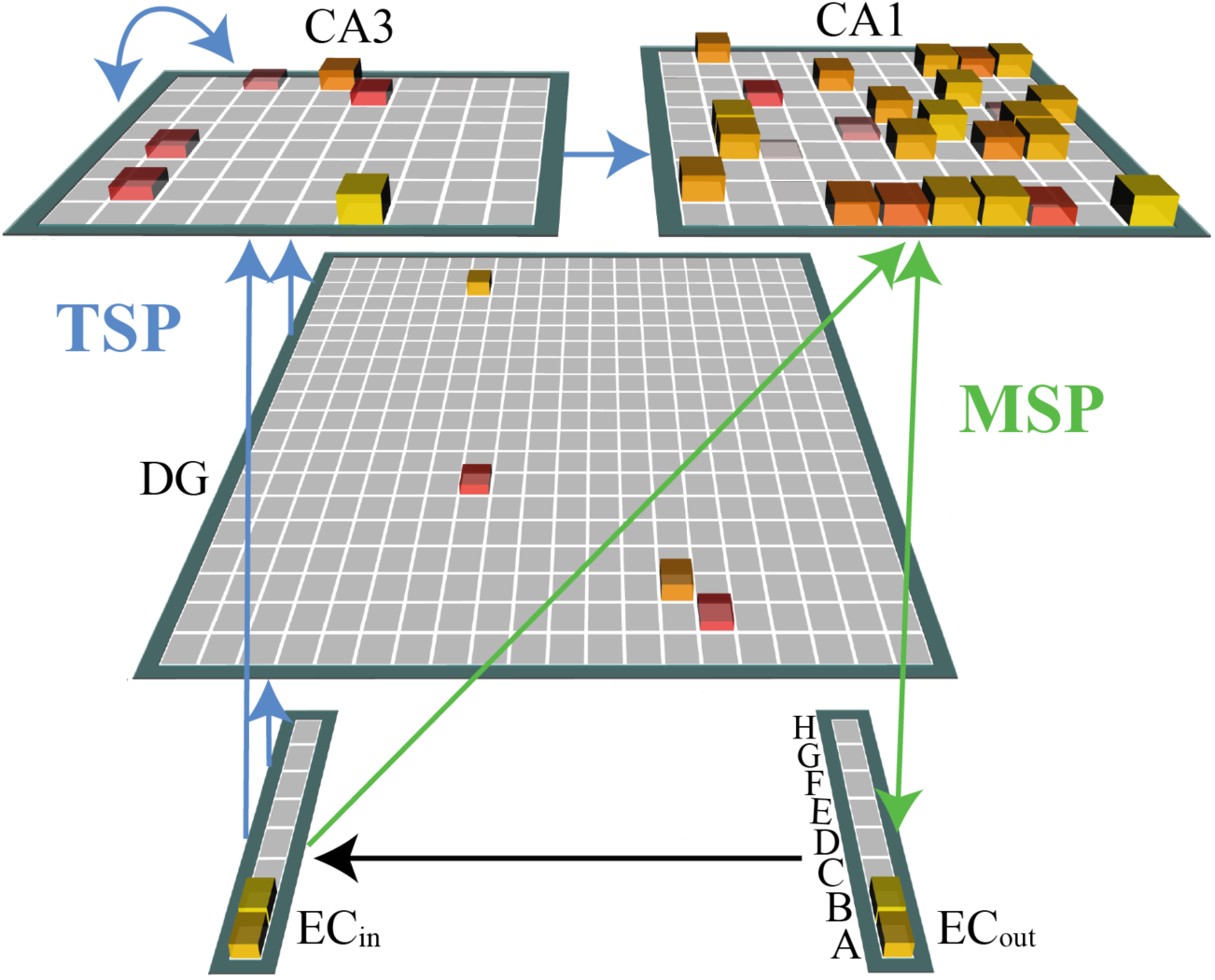
Model architecture. EC_in_ serves as input and EC_out_ as output for the network. The network is trained to reproduce the pattern of activity in EC_in_ on EC_out_. Three hidden layers — DG, CA3, and CA1 — learn representations to support this mapping, with activity flow governed by the projections indicated by the arrows. Blue arrows make up the TSP and green arrows make up the MSP. This snapshot shows network activity during pair structure learning, where pair *AB* is presented to the network and successfully reproduced in EC_out_. The height and yellowness of a unit both index its activity level.

#### TSP

EC_in_ projects to DG and CA3 in the TSP. These projections are sparse: each DG and CA3 unit receives input from 25% of the EC_in_ layer, and the “mossy fiber” projection from DG to CA3 is even sparser (5%). In addition, DG and CA3 have high levels of lateral inhibition, which further limit the amount of activity. These features allow DG and CA3 to avoid interference by forming separated, conjunctive representations of incoming patterns, even when the patterns are highly similar. CA3 also has a fully connected (every unit to every other) projection to itself, which helps bind pieces of a representation to one another and retrieve a full pattern from a partial cue. CA3 then has a fully connected projection to CA1, completing the TSP.

#### MSP

There are fully connected projections in the MSP from EC_in_ to CA1, CA1 to EC_out_, and EC_out_ to CA1. CA1 has much less local inhibition than DG and CA3. These two properties — full connectivity and low inhibition — encourage overlapping representations in CA1 for patterns with similar inputs and outputs. See Supplementary Material for parameter details (Tables S1-2), and for discussion of the removal of the MSP “slots” used in previous versions of the model.

#### Network initialization

For each simulation, we ran 500 networks. Each network corresponds to a particular randomized configuration of the sparse projections in the TSP, and to randomly reinitialized weights throughout the network. These re-initializations were treated as random effects in statistical tests.

### Learning

The model is trained to adjust connection weights between units such that it can duplicate the patterns presented to EC_in_ on EC_out_. The learning in the version of the model used here (13) is different from older versions in two important ways. First, whereas previous versions used only Hebbian learning, this version uses a combination of error-driven and Hebbian learning (12, 13, 19). The error-driven component uses Contrastive Hebbian Learning (20), which adjusts connection weights such that activity during a “minus phase” becomes more similar to activity during a “plus phase.” Incorporating error-driven learning into the model results in increased memory capacity (13). Second, whereas previous versions pre-trained the MSP, this version trains the MSP online, at the same time as the TSP, which also results in better memory capacity (13).

The learning procedure used here is based, as in other models (21), on empirical findings of differences in projection strengths between subfields at different phases of the hippocampal theta oscillation (22). At the trough of the theta cycle, EC has a stronger influence on CA1, whereas at the peak, CA3 has a stronger influence on CA1. The model instantiates this as two minus phases on each trial. In one, EC_in_ projects strongly to CA1, and CA3 → CA1 is inhibited; this corresponds to a discretized sample of the theta trough. In the other, CA3 projects strongly to CA1, and EC_in_ → CA1 is inhibited; this corresponds to a discretized sample of the theta peak. These phases have been proposed to correspond to encoding-and retrieval-like states: A strong influence of EC_in_ on CA1 — a state where the external environment directly influences CA1 — is considered more akin to encoding, while a strong influence of CA3 on CA1 is more akin to retrieval (21).

Activity during both minus phases in a trial is contrasted with activity during a plus phase, in which the target pattern is clamped on EC_out_. Weights are changed such that patterns of unit coactivity during the minus phases are shifted more toward those of the plus phase, which promotes better mapping of the pattern in EC_in_ to EC_out_. Modification of the model’s internal representations to better align with the observed environment is a general property of error-driven learning algorithms, but in the case of the hippocampus may be related to the idea that the region carries out match/mismatch computations (23, 24). The learning rate in the TSP is 10x higher than in the MSP. See (13) and Table S2 for more details. Note, however, that we use the version of the model implemented in (18), which uses Contrastive Hebbian Learning for all learnable pathways, as opposed to pure Hebbian learning used for some TSP projections in (13).

### Stimulus presentation

To simulate sequential item presentation, we presented items to EC_in_ using a moving window that encompassed the two most-recently presented items; this approach is based on findings that the representation of a previous stimulus persists over delays in EC (e.g., 25). We included temporal asymmetry, presenting the current stimulus with full activity (clamped value = 1) and the previous stimulus with decayed activity (0.9).

### Testing and analyses

We tested the network with trials in which each of the stimuli was presented by itself. On each trial, we clamped the activity of the Input layer unit for the stimulus to 1 and recorded the activity level of each unit throughout the network after 20 cycles of processing (initial response), and after the network had fully settled (we used 80 cycles, though 40-50 cycles are usually enough). We chose 20 cycles for the initial response because at that point activity has spread throughout the network, including to EC_out_, but not from EC_out_ to EC_in_ (preventing big-loop recurrence). For some simulations, we also looked at the activity evoked by pairs of items, in which case input units for both items were clamped to 1.

During test, network representations were assessed in a neutral state, where the projections between layers were scaled to values in between the encoding-like and retrieval-like states described in *Learning*. No connection weights were changed during testing. Networks were tested before training (epoch 0) and after every epoch of training.

#### Output analyses

We assessed the probability, after settling, of activating a particular item in EC_out_ above 0.5 given presentation of a particular item in EC_in_.

#### Pattern analyses

We recorded the pattern of unit activity evoked by each test item, in each hidden layer, for the initial and settled response. We calculated Pearson correlations between the patterns evoked by different items.

All analyses were done within each of 500 randomly initialized networks, and results were then averaged across networks.

### Lesions

To lesion the TSP, we set the strength of the following pathways to 0: EC_in_ → DG, EC_in_ → CA3, DG → CA3, CA3 → CA3, and CA3 → CA1 (all blue pathways in Fig. 1). To lesion the MSP, we set the strength of EC_in_ → CA1 to 0. We did not alter the CA1 → EC_out_ pathway, though it could be considered to be part of the MSP, because this projection is required for producing output in the model. The pathways were lesioned during both training and testing.

## Results

### Learning episodes versus regularities

We tested whether a model of the hippocampus designed to simulate episodic memory (12, 13) can pick up on statistics in continuous sequences. In prior simulations using this model, the episodes to be learned were clearly demarcated for the model; for example, the model was used to simulate experiments in which subjects were shown word pairs and asked to memorize that they go together (12). By contrast, in statistical learning experiments, there is a continuous sequence of stimuli with no demarcations of event boundaries. The only way to detect the regularities is to track statistics across experiences.

We exposed the model to sequences containing embedded pairs. There were eight items (*A*-*H*) grouped into four pairs (*AB*, *CD*, *EF*, *GH*). Items within a pair always occurred in a fixed order but the sequence of pairs was random. Specifically, the second item in a pair could transition to the first item in one of the three other pairs. Back-to-back repetitions of a pair were excluded, since this is a common constraint in statistical learning experiments and because allowing repetitions would dilute the temporal asymmetry (both *AB* and *BA* would be exposed). There was a moving window of two stimuli presented at a time. After *AB*, for example, *BC, BE*, or *BG* followed with equal probability; if *BC* was chosen, the next input would be *CD*. To detect regularities, the model had to be sensitive to the fact that, over time, pairs (e.g., *AB*) occurred more often than sets of two items spanning pairs (e.g., *BC*). To contrast this learning challenge with a more “episodic” situation with demarcated events, we also ran simulations where pairs were presented in interleaved order, with no transitions between pairs. In other words, *AB*, *CD*, *EF*, and *GH* all appeared but never *BC* or *FG*, for example. Note that we will use *A* and *B* throughout the results to refer to the first member and second member of a pair, respectively. All results are averaged across the four pairs.

Different representations emerged in the model for sequences that did vs. did not require sensitivity to statistics across trials. For the latter, “episodic” sequences, the network quickly learned to activate both members of a pair in EC_out_ (*A* and *B*), when presented with either pairmate individually (*A* or *B*; Fig. 2C). The hidden layers learned representations that allowed the model to perform this mapping between EC_in_ and EC_out_ (Fig. 2A). In particular, CA3 and DG in the TSP rapidly memorized conjunctive representations of each pair (i.e., memorized the distinct pattern of activity evoked by presentation of both pair members at once). Because each item had only been viewed in one pair, this memorization caused each pairmate, on a given test trial, to immediately pattern complete to this conjunctive representation. CA3’s initial pair similarity was somewhat lower than DG’s because the two sets of sparse projections — EC_in_ → DG, DG → CA3 — make it more difficult for CA3 to integrate across paired items.

**Figure 2.**
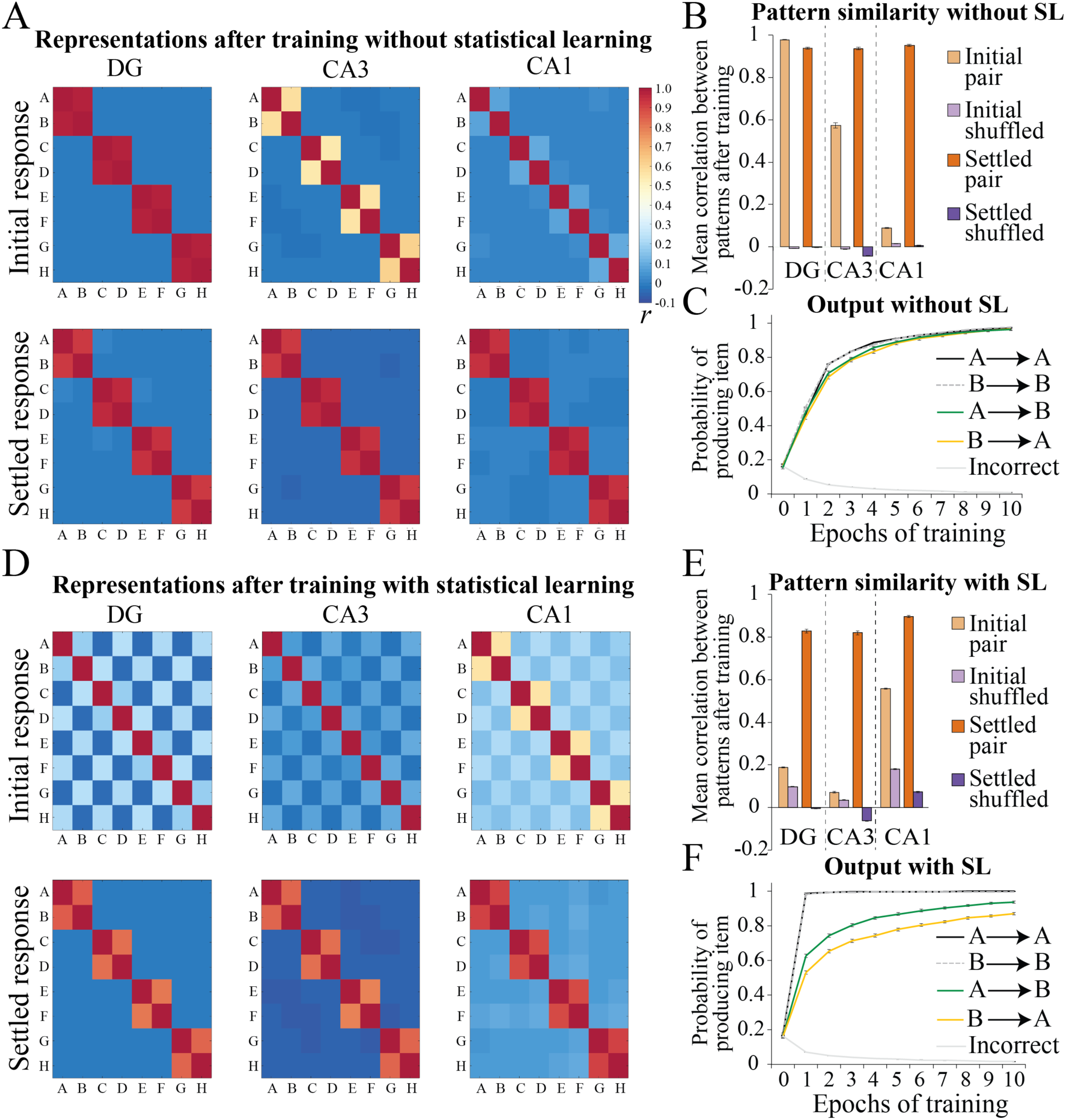
Pair structure. **(A)** Average representational similarity across networks in each of the three hidden layers of the model, after training on episodic sequences that did not require statistical learning (SL). In the heatmaps, each of the 8 test items appears in the rows and columns, the diagonals correspond to patterns correlated with themselves, and the off-diagonals are symmetric. **(B)** Average representational similarity by pair type, for initial and settled response. “Shuffled” pairs are items paired with all other items that were not the trained pairmate (*AC*, *AD*, etc.), including both viewed and unviewed pairings. **(C)** Average probability of activating a particular item on the output given a particular item on the input, over training. For example, *A* → *B* is the probability of activating the second member of a pair above threshold given the first. “Incorrect” is the probability of producing an item that is not the current item or its pairmate. Each input was presented once per epoch in permuted order. **(D, E, F)** Same as above, for sequences that required SL. Each pair was presented approximately 5 times per epoch (with 80 total inputs per epoch). For all subplots, values are means across 500 random network initializations. Error bars denote +/– 1 SEM across network initializations. Some error bars are too small to be visible.

Initial similarity in CA1 was much weaker (Fig. 2B) simply because CA1 learns more slowly (CA1 initial pattern similarity becomes higher after longer training, not shown), but this slower learning rate does not detract from CA1’s ability to help communicate information in CA3 to EC_out_. After the pattern is completed in EC_out_ and information travels to EC_in_, the similarity structure is then strongly apparent in all hidden layers.

What happens when the events to be memorized are not demarcated and must be learned over time from temporal statistics? The model was still able to learn the pairs, activating *B* on EC_out_ given *A* on EC_in_ and vice versa (Fig. 2F), despite each of those items also being exposed with three other items. Unlike the previous simulation, there was a striking lack of pair-related similarity in the initial response in DG and CA3 after training (Fig. 2D). Pair similarity was slightly higher than shuffled pair similarity (Fig. 2E) but not for the subset of shuffled pairs that appeared as across-pair transitions (e.g., *DA*; Fig. S1).

The checkerboard structure in DG and CA3 reflects the fact that both *AB* and *DA* occurred in the sequence, but never *AC*. That is, these regions are sensitive to which items have co-occurred, but do not preferentially retrieve an item’s most frequently co-occurring mate. DG memorizes every exposed input pattern, and when given a cue like *A* that is part of more than one pattern, activates units involved in all conjunctive representations of exposed patterns that included *A* (including *AB* but also the other exposed patterns *DA*, *FA*, and *HA*). After the network retrieves *B* on EC_out_ (due to the influence of CA1, as described below), *B* in addition to *A* becomes active in EC_in_, and the DG representation shrinks down to only the units involved in the *AB* conjunctive representation. Pair structure emerges after the network has settled for this reason. The dynamics are similar though not quite as clean in CA3.

In contrast to the previous simulation, CA1 here showed robust similarity for the paired items in its initial (and settled) response. This is because there was enough exposure for CA1 to learn representations with its slower learning rate, and because distributed, overlapping representations are highly sensitive to frequency statistics over time (26).

These simulations indicate that the TSP and MSP form different representations depending on the learning problem. When sensitivity to statistics is not required, the TSP memorizes each of the presented patterns and can complete them from a partial cue, while CA1 simply acts as a translator between CA3 and EC. When sensitivity to statistics *is* required, the TSP unhelpfully captures both the pairs and the transitions between them, while the MSP is able to represent the frequencies of pair co-occurrence. In both cases, pattern completion expressed in EC_out_ causes the pairmate to arrive on EC_in_, and the pair’s representation activates throughout the network.

### Representational change over time

Though the TSP does not represent regularities in the initial response by the end of training, there is an earlier period where it does represent them weakly and better than the MSP (Fig. S1A). This occurs because the TSP rapidly learns whatever it is exposed to, and the true pairs *AB* do occur more frequently. However, as the network gains more exposure to all transitions (*BC*, *DA*, etc.), weights max out and the similarity for across-pair transition fully catches up. As *B* pulls away from *AB* and toward *BC*, the relatively greater similarity for pairmates weakens. CA1’s similarity structure does not weaken over time in this way, even with extensive training (Fig. S1C-D). To understand the impact of these TSP dynamics on the model’s behavior, we ran simulations with the MSP lesioned. The TSP alone initially supported weak retrieval of the correct pairmate, but over time, real within-and across-pair transitions were retrieved equally (Fig. S1B). Thus, by the end of training, a network with an MSP lesion exhibits virtually no sensitivity to regularities.

### Temporal asymmetry

Because items were presented two at a time, with the previous item less active than the current item, we can examine temporal asymmetry in the network. The hippocampus is known to be involved in prediction (6, 7, 27–29) and so the network should show a bias for temporally forward vs. backward pattern completion. This was true in the network’s output: *B* was more activated in the output layer by *A* input than vice versa (Fig. 2F). This results from the higher activity of *B* compared to *A* when *AB* is presented in training, causing more strengthening of the weights connecting to *B*’s output unit than to *A*’s. There was asymmetry in the hidden layer representations as well: The settled pattern of activity evoked by *A* alone was more similar than *B* alone to the settled conjunctive *AB* pattern (DG: *r* = .94 vs. .87, respectively, t[499]=8.10, p<0.001; CA3: .93 vs. .86, t[499]=8.01, p<0.001; CA1: .96 vs. .93, t[499]=7.82, p<0.001).

### Statistical learning with an undeveloped TSP

Infants are remarkably good at statistical learning (30, 31), and our model suggests that this may be because the MSP develops earlier than the TSP (32, 33). To test this idea, we lesioned the TSP. This simulates, for demonstration purposes, the boundary case of having no ability to use that pathway, though infants may have some limited use. We found that learning and representations were essentially unchanged with a lesioned TSP (Fig. S2). In fact, pairmate retrieval was slightly better without vs. with the TSP (at the end of training, mean *A* → *B* retrieval: .98 vs. .94, respectively, t[998]=6.66, p<0.001; mean *B* → *A* retrieval: .90 vs. .87, t[998]=2.49, p=0.01). This stands in contrast to the very detrimental effects of an MSP lesion (Fig. S1B), revealing that the MSP is necessary and sufficient for statistical learning and providing a possible explanation for intact rapid statistical learning in infants despite protracted hippocampal development.

### Higher-level learning

The structure of the sequences considered so far was simple — it could be learned by tracking the strength of transition probabilities between adjacent items or the joint frequency of pairs. To explore whether higher-level statistics can be learned in the MSP, we simulated learning of a “community structure” sequence that cannot be parsed based on transition probability or joint frequency (5, 34). The sequence was generated via a random walk on a graph (Fig. 3B) with three densely interconnected “communities” of nodes (35). The walk tended to stay in a community for awhile before transitioning to the next, but any individual node had an equal probability of transitioning to exactly four other nodes. We constrained the sequences such that the observed joint frequencies were exactly equated in each epoch (as opposed to equated on average). Thus, transition probabilities did not provide information as to the location of community boundaries.

**Figure 3.**
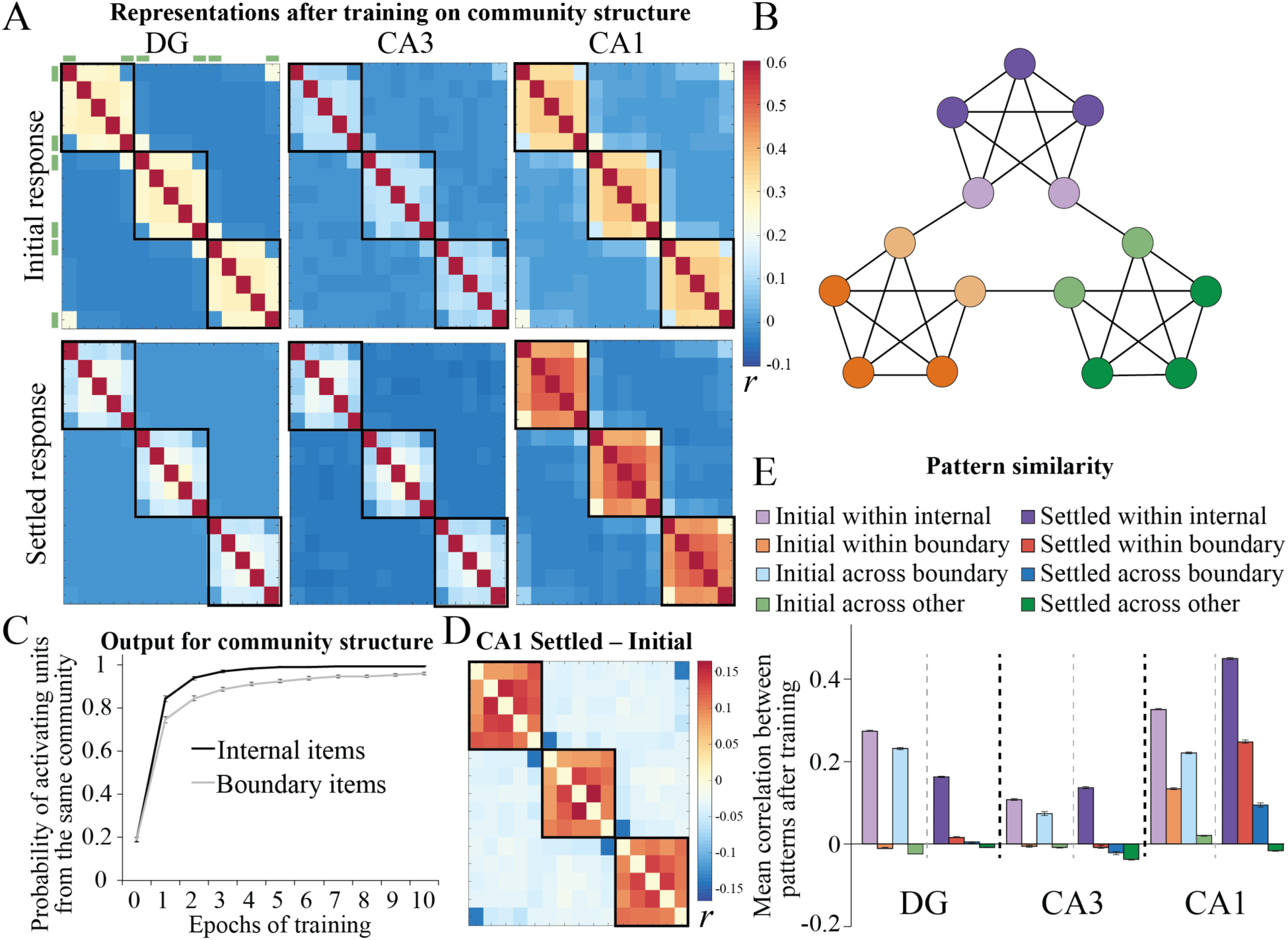
Community structure. **(A)** Average representational similarity after training, with items arranged by community (black boxes; green bars mark boundary nodes). **(B)** Each node on the graph represents a particular item, and the edges indicate which transitions were allowed. **(C)** Average probability of activating units from the same community given an internal item (dark-shaded node) or boundary item (light-shaded node) as input, over the course of training. **(D)** Difference between the settled and initial heatmaps in CA1. **(E)** Average representational similarity between two internal nodes from the same community (within internal), the two boundary nodes from the same community (within boundary), two adjacent boundary nodes from different communities (across boundary), and all other pairs of items from different communities (across other).

The only change to the architecture and parameters for this simulation was the addition of 7 units to EC_in_ and EC_out_ to accommodate each of the 15 nodes in the graph. We exposed the model to sequences in the same way, with two items presented at a time and the previous item less active than the current item. (Temporal asymmetry is not meaningful here, though, as every pair occurs in both orders.) There were 60 inputs per epoch, presented over 10 epochs. Because all pairs occurred with equal frequency, to learn the communities, the model had to pick up on the fact that items in the same community appeared with overlapping sets of other pairmates, whereas items from different communities did not.

The model successfully learned to activate other items from a test item’s community (Fig. 3C). Notably, this was true considering only the boundary items: By the end of training, tested boundary items almost never activated the adjacent boundary node in a different cluster over one of the adjacent nodes in the same cluster. The network again exhibited different representational similarity in the initial and the settled response (Fig. 3A): DG and CA3 initially represented all exposed pairs but largely ignored higher-level structure; in contrast, CA1 was immediately sensitive to higher-level structure. This is best illustrated at community boundaries: the two boundary nodes in the same community were never seen together in training, whereas two adjacent boundary nodes from different communities were. Thus, high similarity for within-community boundaries means that a region has learned structure beyond basic co-occurrence statistics. This was found in the initial response in CA1 (Fig. 3E), although it also had high similarity for across-community boundaries. DG and CA3 showed low similarity for within-community boundaries and high similarity for across-community boundaries, indicating that these regions did not learn the higher-level structure.

After allowing big-loop recurrence, the higher-level structure became clearer in CA1, and throughout the network. Community structure fully overtook temporal proximity, with an increase in within-community boundary similarity and a decrease in across-community boundary similarity (Fig. 3D-E). To understand why, consider what happens when a boundary item is presented at test: it retrieves a directly linked associate, which is more likely to be an internal node from the same community than the boundary node from the other community, because of greater within-community internal vs. across-community boundary similarity (Fig. 3E); EC_out_ will send activity for that internal node to EC_in_, which then travels to the hidden layers and further emphasizes the representation of that community. We verified that the difference between the initial and settled representational similarity was indeed attributable to big-loop recurrence (vs. recurrence in activity between CA1 and EC_out_), by lesioning either EC_out_ → EC_in_ (needed for big-loop recurrence) or EC_out_ → CA1 during testing. Higher-level behavior was unchanged in the EC_out_ → CA1 lesioned network but much weaker in the EC_out_ → EC_in_ lesioned network.

### Associative inference

Learning the higher-level structure in the community structure simulations required transitive associations — associations that are not based directly on observed item pairings but rather on the fact that different pairs *share* associates. Other learning paradigms, such as transitive inference, acquired equivalence, and associative inference, have the same property, though they are not considered conventional statistical learning paradigms in that there is not a continuous presentation of individual items. Nevertheless, these paradigms all involve hippocampal processing and, like statistical learning, require integration across experiences (36).

To demonstrate the close relationship between our simulations of statistical learning and these paradigms, we present a simulation of associative inference. In this paradigm, two items *A* and *B* are presented together, and in separate trials, *B* and *C* are presented together. Humans and animals are then able to infer that *A* goes with *C* (36). We trained the model on three triplets with this structure: *ABC, DEF*, and *GHI*. We presented each pair 10 times per epoch in permuted order, over 20 epochs. Because the paradigm involves presenting two stimuli at a time, both stimuli in an input were presented with full activity. We made one change to the network parameters in addition to setting the number of units in EC_in_ and EC_out_ to 9. To allow the model the possibility of activating the transitive associate (*C* given *A*) and not just itself and its direct associate (*A* and *B* given *A*), which is its strong tendency with *k=*2, we lowered inhibition to *k*=3 when testing the network (during training, *k*=2, as before). This is critical for accounting for transitive behavior. See Figures S3-4 for discussion of this point and simulations showing how *k*=3 at test affects pair structure and community structure simulations (to summarize, it enhances higher-level structure in the latter and adds a small amount of noise to the former).

The results of these simulations matched the community structure simulations. In the initial response, DG/CA3 represented direct pairs *AB* and *BC*, but not transitive pair *AC*, whereas CA1 had graded similarity structure that reflected transitive relationships (Fig. 4A).

**Figure 4.**
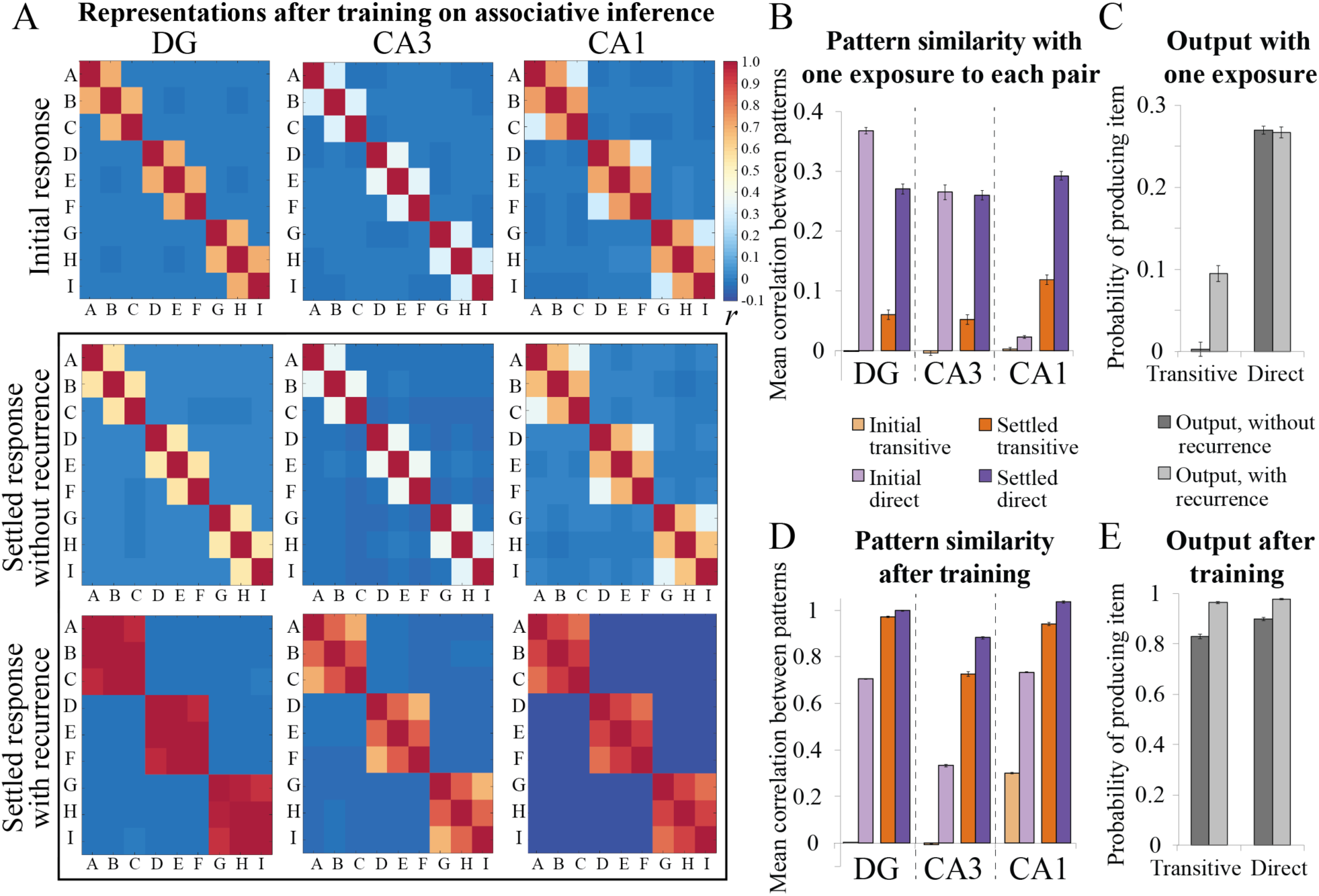
Associative inference. **(A)** Average representational similarity after training, with items arranged by triplet. The settled response is shown with and without recurrence allowed. The initial response was very similar for the two variants and is shown with recurrence. **(B)** Similarity structure after one training trial with each of the direct pairs. The correlation between direct *AB* pairs and between transitive *AC* pairs is shown, subtracting the correlation for shuffled pairs as a baseline. This is shown for the initial and settled response, both in networks with recurrence (though only the settled response is affected by recurrence). **(C)** The probability of producing the direct pairmate (*B* given *A*) and transitive pairmate (*C* given *A*), subtracting the probability of producing other items as a baseline, with and without recurrence. We allowed any above-zero activity in EC_out_ units to count as “producing the item” to simulate the sensitive forced choice test used in associative inference (14). **(D, E)** Same as **B, C** for fully trained network.

There has been recent debate as to whether this task is solved using recurrence or “static” representational similarity — overlap in representational space that is not a result of recurrent activity dynamics (15, 36–38). To evaluate this in our model, we contrasted pattern similarity in the settled response for networks with and without recurrence during test. Networks without recurrence had EC_out_ → EC_in_ and EC_out_ → CA1 connections lesioned (only in the test phase), so that representational similarity and behavior must be caused by the representations in the initial response. The transitive structure was present without recurrence in CA1, but became stronger throughout the network with recurrence (Fig. 4A), indicating that both static representational similarity and recurrence contribute to transitive associations in the model.

Another way to evaluate the role of recurrence is to assess representations and behavior very early in training, before CA1 has time to develop overlapping representations for transitive associations. After just one exposure to each of the direct pairs, when there was no transitive structure at all in the initial response in CA1 (Fig. 4B), recurrence could support transitive behavior (Fig. 4C). Indeed, recurrence was *necessary* for this behavior, as the network without recurrence could produce direct pair but not transitive output after one exposure. The network can use rapidly memorized associations between the direct pairs in the TSP in addition to recurrent dynamics to link from *A* to *C* through *B*. After recurrent activity settles, the transitive associate appears in EC_in_, allowing all hidden layers to show apparent transitive pattern similarity (Fig. 4B). CA1 notably showed stronger settled similarity here than DG and CA3 because of its lower sparsity.

After full training, CA1 showed transitive similarity in the initial response (Fig. 4A,D). This representation was sufficient to support transitive behavior, as transitive associates were produced in EC_out_ even when recurrence was not allowed (Fig. 4E). Allowing recurrence enhanced the similarity structure throughout the network (Fig. 4A,D) as well as the transitive behavior (Fig. 4E).

## Discussion

Avoiding interference between related memories is often critical, but it can be just as important to notice their commonalities. Our simulations suggest that there may be complementary learning systems within the hippocampus that subserve these competing goals — a microcosm of the broader CLS theory of hippocampus and cortex. We found that DG and CA3, forming the TSP to area CA1, robustly represented distinct episodes but failed to learn regularities across episodes. This is due to aggressive pattern separation of similar experiences in these regions, caused by sparse connectivity and high inhibition. In contrast, the direct MSP pathway to CA1 learned regularities across experiences. This is facilitated by overlapping representations in CA1 that result from full connectivity, lower inhibition, and a slower learning rate. The MSP can function even when the TSP is fully lesioned, which might provide an explanation for why infants, who have undeveloped TSPs, are prodigious statistical learners. The overlapping representations in CA1 are akin to “nodal codings” that have been observed there, with shared features across events represented by overlapping populations of neurons (39, 40, see also 41). These representations have been proposed to support relational memory and generalization (42), consistent with our account.

The community structure and associative inference simulations demonstrated that the model is also capable of learning transitive associations. For community structure, CA1 learned to represent the two boundary nodes in the same community more similarly despite the fact that they were never experienced together. This was possible because they shared overlapping sets of associates — a principle we previously demonstrated in a simpler neural network model of temporal community structure (34). More generally, our model predicts that static representational similarity will emerge in CA1 whenever there are correlations in EC inputs and/or targets, and will reflect input frequencies. These principles of learning mirror those demonstrated in many prior neural network models that use error-driven learning and overlapping, distributed representations (e.g., 26, 43).

Our simulations are consistent with findings from recent human fMRI studies that used pattern similarity to assess changes in representations across voxels in the hippocampus. In a high-resolution study, we found that neural representations of items that were consistently paired in a continuous sequence became more similar in all subfields of the hippocampus (6), analogous to the settled response in our pair structure simulations. Similarly, in a study of temporal community structure, we found that hippocampal representations of items from the same community became more similar than those from different communities (5). Although this study was not performed at high resolution, probabilistic segmentation suggested that the effect was most reliable in CA1, consistent with the stronger CA1 effects in our simulations. Finally, an associative inference study found the strongest pattern similarity for transitively associated pairs in CA1 (14); direct pairs were presented only once in this study, corresponding to our simulations using one exposure (Fig. 4B).

Our model builds on two recent neural network models of the hippocampus. The first is Ketz et al. (2013), who developed a technique for training the MSP online with error-driven learning. This allowed us to explore what kinds of representations emerge during training in the MSP, which was not possible in earlier versions of these models. Earlier models posited that MSP connections did not show appreciable (or any) learning on the timescale of a single experiment (44), but there is now ample evidence that the MSP does learn at this timescale (45–47). The second is REMERGE (15), which demonstrated the utility of big-loop recurrence in transitive inference, paired associate inference, and acquired equivalence. Such recurrence plays three roles in our simulations: (i) it allows the structure learned in CA1 to spread to DG and CA3; (ii) it strengthens the similarity structure in CA1, due to the actual presence of the pairmate representation in EC_in_; and (iii) when pairwise co-occurrence is not sufficient to uncover structure (i.e., community structure and associative inference), big-loop recurrence enhances transitive relationships. Like REMERGE, our model learns statistics while protecting the role of pattern-separated representations in episodic memory. In contrast, however, our model outlines a role for static representational similarity in the hippocampus.

### Static representations versus recurrence

There is a debate in the hippocampal generalization literature as to whether transitive associations are supported by overlapping memory representations formed during encoding or by recurrent activity online at retrieval (15, 36–38). The core question is whether generalizations are stored in weights in the network or dynamically computed as needed. Our account is a hybrid between REMERGE, which depends entirely on recurrent dynamics, and models that depend entirely on overlap in pair representations (37, 48). In our model, recurrent dynamics play a dominant role early in training, before the MSP has a chance to learn overlapping representations (this dependence on number of exposures was also proposed by 36). After sufficient exposure, however, both static representational similarity and recurrence contribute.

Evidence for both mechanisms exists in the human fMRI literature (36). For example, in acquired equivalence, increased hippocampal activity over training that correlates with generalization performance has been taken as evidence of encoding of integrated representations (49), and in associative inference, higher hippocampal activity for transitive pairs at test has been interpreted as flexible recombination of direct pairs at retrieval (50). However, REMERGE accounts for the acquired equivalence results using only recurrent dynamics (15), and our model can produce transitive associative inferences at test with static representational similarity (Fig. 4D-E).

How, then, can these theories be disentangled? One way is that our model and REMERGE make different behavioral predictions for blocked vs. interleaved stimulus presentations. In blocked presentations, a set of inputs X is fully learned before moving on to learning a second set of inputs Y, whereas in interleaved presentation, inputs from X and Y are interspersed. Networks with overlapping representations show effective learning with interleaved training but catastrophic interference after blocked training, where X is forgotten during learning of Y (1). In our model, the use of overlapping representations in CA1 would be expected to lead to catastrophic interference in blocked learning designs, whereas the orthogonalized representations used in REMERGE would be relatively unaffected. Correspondingly, our model but not REMERGE predicts a benefit for interleaved stimulus presentation on hippocampal generalization, after CA1 has had enough exposure to develop overlapping representations. Behavioral experiments in which learning occurs on a timescale that is likely to be hippocampally dependent suggest a benefit for interleaved learning (e.g., 51, 52), though under some conditions blocked learning can be beneficial, which neither model predicts (53).

Our model also makes specific predictions that could be tested in electrophysiological experiments using population recordings at different sites in the hippocampus. After learning temporal regularities, the pattern of activity evoked by a single item should result, over the course of one trial, in a pattern of activity similar to temporal associates first in CA1, then in deep layers of EC, then superficial layers of EC, then in DG and CA3. Lesions to any portion of the TSP should not affect behavioral evidence of statistical learning, nor should it affect representations in CA1. If lesioning only the projection from CA3 to CA1, all representational effects should be preserved and appear in the order just described. Conversely, lesions to the MSP should weaken representational and behavioral effects. Our model makes the counterintuitive prediction that an animal with a specific MSP pathway lesion should be weakly sensitive to pair structure at first, but that this ability should then deteriorate over time. REMERGE does not contain hippocampal subfields and so cannot address these predictions.

### Anterior versus posterior hippocampal function

There is an anatomical gradient in the prominence of different hippocampal subfields, in which regions CA1-3 are relatively overrepresented in anterior hippocampus and DG is over-represented in posterior hippocampus (54). Considering DG as an important contributor to pattern separation in the TSP, this suggests there may be a relatively more dominant role of the MSP in the anterior hippocampus. Our model then makes the prediction that the anterior hippocampus should be more dominant in statistical learning. This is consistent with human fMRI findings on the anterior hippocampus: (i) it generally shows stronger effects in pair structure and community structure paradigms (5, 6); (ii) it plays a stronger role in integrating memories in the associative inference task (50, 55); (iii) it fully integrates across elements in an event narrative analogue of the associative inference task (56); (iv) it is active during transitive inference (57); and (v) its activity varies with generalization performance in acquired equivalence (49). This generalization of experiences across time in anterior hippocampus may be related to the gradient of spatial representations found in rodent hippocampus, where ventral (anterior) place cells represent larger, more overlapping areas of space compared to dorsal (posterior) ones (56, 58, 59). Analogous to our account, this varying spatial scale has been proposed to allow the hippocampus to carry out the complementary tasks of generalizing and avoiding interference in spatial memory (60).

### Temporal and sequence processing

There is substantial evidence from the rodent literature that the MSP and CA1 play a special role in temporal processing, including that: inhibition of the MSP leads to deficits in temporal association memory (61), CA1 supports temporal order memory (62–64), hippocampal place fields in CA1 expand to represent earlier positions on a track (65), and CA1 can generate reliable sequential spiking patterns on its own (66). There is also evidence from human fMRI that CA1 plays a larger role than other hippocampal subfields in sequential processing (34, 67).

These findings are consistent with our account of the MSP’s role in extracting temporal regularities across experiences. They are surprising, however, given the large hippocampal modeling literature that has focused on the role of recurrent heteroassociative connections in CA3 in processing temporal sequences (24, 68–72). (See Rolls and Kesner (73), though, for discussion of the consonant idea that less sparse representations in CA1 may allow it to represent sequences better than CA3.) One possibility is that greater reliance on the TSP vs. MSP in sequence processing may depend on the type of sequence. If there is an obvious repeating sequence, the TSP may be best suited to learning and reproducing it, whereas a noisier sequence that requires integration over time may require the MSP. It is also possible that big-loop recurrence or recurrence between EC and CA1 may play a similar computational role to recurrence within CA3. In our model (as in REMERGE), input *A* retrieves associated item *B*, which then travels through the big loop, such that the model first represents *A*, then additionally *B*. We tried training the model on a longer sequence (data not reported here), *ABCDEFGH*, and found that when given *A* at test, and with some synaptic depression, it would use the big loop to sequentially retrieve *B* (as *A* fades due to depression), then *C* (as *B* fades), then *D*, etc. This indicates that big-loop recurrence can be sufficient to produce robust sequential behavior.

### Persistence of item activity over time

We chose to represent two items at a time in EC based on findings that it exhibits persistent activity for recently-exposed stimuli (25, 37). Indeed, lesions to parahippocampal regions but not to hippocampus proper impair delayed non-match to sample performance, suggesting that these regions may be more important than the hippocampus in maintaining simple traces of past stimuli (74–76). For purposes of demonstrating model learning principles, we adopted the simplified approach of allowing only the immediately preceding item’s activity to persist for the presentation of the current item. Depending on the speed of item presentation and the specifics of the paradigm, however, there could be longer histories of item persistence (77). How this would impact model representations and behavior is an interesting area for future work.

### MTL cortex and beyond

A related division of labor between episodic memory and statistical learning in the MTL has previously been proposed, with the hippocampus proper supporting episodic memory and EC learning incrementally and coming to represent co-occurring stimuli more similarly (78, 79). Empirical support for this hippocampus-EC dissociation comes from animal studies of incremental conditioning over days (80, 81). Although we did not model learning within EC, we expect that it will learn overlapping representations as in CA1, just on a slower timescale of days, not minutes. A slower learning rate in MTL cortex would still be consistent with it supporting rapid (even one-shot) familiarity for individual items (4, 12), which can be instantiated as subtle changes to the representation of one item, as opposed to the more computationally difficult binding across multiple items. It would also be consistent with findings of rapid learning-related changes in representational similarity in MTL and other cortical areas (6). This is because the hippocampus can reinstate an associated item in MTL simultaneously with processing of the current item in the same areas, causing apparent representational similarity without local learning.

One intriguing possibility is that there is a hierarchy of learning rates, perhaps related to a hierarchy of temporal window sizes for information accumulation (82), where the TSP is fastest, followed by the MSP, then MTL cortex, then cortical areas like the anterior temporal lobe that integrate across different types of sensory and motor information (83), and then finally areas that support specific sensory or motor functions. There is substantial evidence that the MSP/CA1 operates on a longer timescale than the TSP/CA3 (73, 84–87), consistent with our model’s relatively slower learning rate there. Further down the hierarchy, after days of learning to associate pairs of fractals, perirhinal cortex exhibits representational similarity for paired fractals (88–90). Such “pair coding” neurons are found even further down in inferotemporal cortex (IT), but their response appears about 350 ms later than in perirhinal (88), they are more prevalent in perirhinal cortex (89), and IT pair coding is abolished by perirhinal and entorhinal lesions (91). For the rapid observational statistical learning considered here — on the order of minutes to hours — the learning rate of the MSP may be most suitable. The site of local learning may then expand outward with increasing exposure and opportunities for consolidation.

The idea that CA1 is intermediate in timescale between CA3 and MTL cortex may also relate to its role as a stable translator between CA3 and EC in episodic learning tasks. It is important for those mappings to be stable in order for CA1 to effectively communicate the details rapidly stored in DG and CA3 back to EC. However, this stability need only last while in a particular context (as opposed to over the lifespan, 44).

Another cortical area with notable relevance is the medial prefrontal cortex. This area has been shown to be involved in an understanding or familiarity with an event (92, 93), it plays a role in learning and expressing transitive associations (94–96), and has been implicated in learning temporal community structure (34). This region may be an important consolidation site for structured event knowledge acquired by the hippocampus (97).

### Conclusions and open questions

Where does this work leave us with respect to the original CLS theory, which posits that the hippocampus stores new information and slowly teaches it to cortex to prevent catastrophic interference (1)? We think that this fundamental principle remains unchallenged: It is still critical to use a separate memory store to protect long-term knowledge representations from novel, potentially interfering information, and we still think the hippocampus plays this role. The modification to the CLS framework that we propose is that both novel episodic and novel statistical information are quickly learned in the hippocampus — indeed, both types of information could interfere with cortical representations if learned directly in cortex. We propose that it is not the *entire* hippocampus that stores pattern separated representations for separate recent experiences, but rather that this is the province of the TSP. The MSP then specializes in extracting regularities across recent experiences. Indeed, CA1 is the most cortex-like area of the hippocampus, with relatively more overlapping representations and a slower learning rate (though still sparser and faster than cortex), which are precisely the properties that encourage neural networks to efficiently generalize across experiences. Similarly, DG and CA3 are the areas that truly instantiate the hippocampal learning properties as presented in CLS.

One open question is whether the analogy to CLS consolidation might hold within this microcosm: On this short timescale, does the TSP train the MSP offline to help avoid catastrophic interference in the MSP, which would be useful in cases of blocked stimulus presentation? Or is the MSP’s utility in rapid statistical learning restricted to cases of interleaved presentation? Another important question is how statistical representations in the MSP affect the process of consolidating information to cortex. CA1 neurons replay generalized trajectories in a maze, not just those directly experienced (98), raising the possibility that the representations in CA1 allow a more sophisticated and potentially more useful transfer of information to cortex (for related discussion and simulations, see 15). Statistical learning in the hippocampus may also encourage replay of the most statistically reliable experiences, as opposed to indiscriminate replay of all experiences.

Future work could also explore whether the two pathways can be controlled depending on the *type* of past information that is currently relevant, or anticipated to be relevant in the future. For example, when we are trying to remember the specifics of a recent experience that overlaps with other recent experiences, perhaps activity in the MSP can be strategically suppressed. Future simulations could assess interactions between the pathways in the context of paradigms in which both episodic details and statistical information are salient or relevant to behavior, such as experiments that demonstrate sampling of trial-unique information after learning statistical information across trials (99).

Though there are many questions that remain to be addressed, we hope the model provides a useful step on the path toward understanding how the hippocampus supports memory for both episodes and statistics.

## Additional Information

## Acknowledgments

The authors thank Samuel Ritter, who contributed to this project during a rotation in K.A.N.’s lab, and helpful conversations with Michael Arcaro, Roy Cox, and Marc Howard.

## Authors’ Contributions

A.C.S. conceived the study, built and ran the models, analyzed the data, and wrote a draft of the manuscript. All authors contributed to the design of the study, discussed the results, and edited the manuscript.

